# The Intergenerational Classroom: A Case Study Integrating Undergraduate and Lifelong Learning Curricula for Biology Education

**DOI:** 10.1101/2025.07.23.666467

**Authors:** Julie Earles, Bethany Stanhope, Kristen Robbins, Alex C. Keene

**Author notes:** Address correspondence to Julie Earles at or Alex Keene.

## Abstract

According to the US Census Bureau (2019) there are approximately 95 million people in the US 55 years of age and older, and that number is expected to increase dramatically in the next decade. This growing population results in an increasing need for STEM learning opportunities for older adults. This need is partially met by lifelong learning communities across the country that offer non-degree programs on a broad range of topics. While many of these programs are located on a college campus, there are few opportunities for interaction between lifelong learners and students in degree-oriented programs. To improve access to STEM education for lifelong learners, we generated an intergenerational classroom composed of lifelong learners and undergraduate students. The semester-long course focused on the biography and medical writings of the late neurologist Oliver Sacks. Our analysis revealed a high level of satisfaction from both undergraduate students and lifelong learners. Both groups overwhelmingly found the intergenerational format beneficial to learning. Furthermore, self-assessments revealed that students felt more positively about intergenerational interactions following completion of the course. Overall, this course provides a framework for increasing access to STEM for the growing older adult population and fostering positive intergenerational interactions. This model could be readily implemented across the country given the abundance of lifelong learning programs currently affiliated with colleges and universities.

---

The number of older adults in the United States has increased dramatically in the past 30 years and is projected to continue this rapid increase. There are currently an estimated 95 million people in the US age 55 and older and 53 million of those are 65 or older [1]. By 2030, the US Census Bureau predicts that there will be approximately 154 million people in the US age 45 and older and 73 million of those will be age 65 and older [1]. Because women in the US live longer than men, the majority of American older adults are women. Growth in the number of older people, accompanied by their important and influential roles in all aspects of society including politics, community, and social structures, underscores the importance of continuing adult education.

Close to 75% of adults think of themselves as lifelong learners, and 25% report having taken at least one class in the past year [2]. This has resulted in the rapid growth of lifelong learning communities and programs that provide informal education and are typically utilized by older adults. The Lifelong Learning Institute Directory produced by the Osher Institutes (2018) lists over 400 lifelong learning institutes in the United States.

A movement in lifelong education is the placement of Lifelong Learning Societies on university and college campuses, providing these groups access to university infrastructure and faculty [3]. For example, there are currently over 120 Osher Lifelong Learning Institutes, with endowments supported by philanthropist Bernard Osher, and most are associated with a university [4,5]. Almost all of the major universities in the State University System of Florida have Lifelong Learning Institutes, including FAU, FIU, UF, FSU, UCF, and USF. Many private colleges and universities also have Lifelong Learning Institutes. For example, in Florida, University of Miami, Nova Southeastern, Eckerd College, and Rollins College all have programs. Lifelong Learning Institutes generally provide informal education because there is no degree or set curriculum.

The recent emphasis on expanded STEM access has primarily focused on the K-12 and college levels, but there is a growing appreciation of the need for informal access to science education throughout the lifespan [6]. Unfortunately, despite rapidly increasing numbers of lifelong learners, STEM education is underrepresented in educational programing. For example, the FAU Osher Lifelong Learning Institute in Jupiter offered over 100 classes from Spring 2017 to Winter 2018 but only one class was STEM-related (See Figure 1). Nevertheless, many adult learners and retirees express an interest in science education, including biological and psychological aging, neuroscience, climate change, and medicine. The underrepresentation of STEM learning highlights the need for novel mechanisms for inclusion of STEM-based classes in lifelong learning societies and more generally, improving access to STEM education for older adult learners. The generation of pedagogy and mechanisms to enhance STEM education for older adult learners in an interactive setting would broadly increase STEM education and scientific literacy in our society.

**Figure 1.**
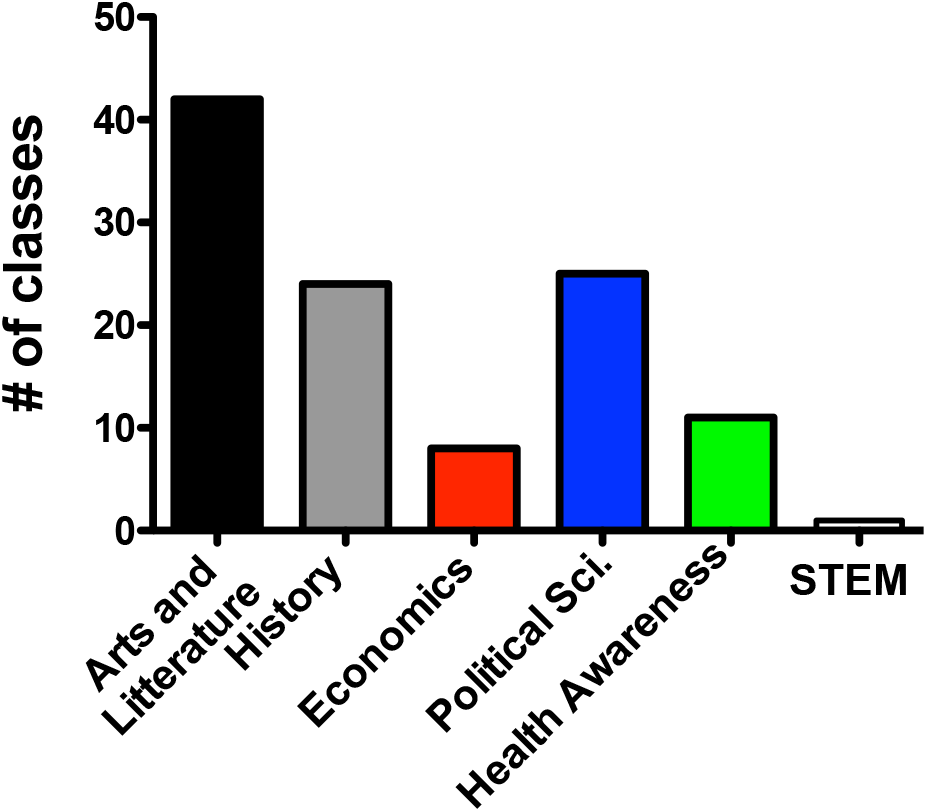
Number of courses/lecture selections by topic at FAU’s Osher Lifelong Institute from Spring 2017 to Winter 2018

To increase the accessibility of STEM education to older adults, and to promote interaction between lifelong learners and undergraduates, we sought to develop and assess the effectiveness of an intergenerational classroom. Despite the physical proximity of most lifelong learning communities to undergraduate students, there is usually little interaction been canonical college programming and the lifelong learning centers. Our intergenerational classroom consisting of both older adults and undergraduate students was designed to capitalize on the benefits of intergenerational learning, including interpersonal benefits, with ages ranging from 20 to 78 years old. Lauren Carstensen, director of the Stanford Center for Longevity, stated “age-related increases in wisdom, life experiences and emotional stability are well-documented, as is a drive to give to others in a meaningful way” [7,8]. There is empirical evidence that older adults often experience an increase in generativity and derive increased satisfaction from helping others [9]. Our intergenerational STEM classroom was designed to allow older adults to help the undergraduate students while at the same time benefiting from the knowledge and ideas of the young students.

Here we describe the development of an upper division, discussion-based neuroscience course integrating Lifelong learners into an undergraduate curriculum. The course focused on integrating the writings of Oliver Sacks, a longtime leader in medical communication, with current topics in neurology and was titled ‘The Life and Science of Oliver Sacks.’ The course was designed to focus on a topic with broad appeal to both undergraduate and Lifelong learning students. The choice to focus on Oliver Sacks was intended to generate interest amongst Lifelong Learners. Oliver Sacks has been a best-selling author and physician dating back to the 1970’s.

The course was also designed to meet a high demand for upper division biology electives due to a large pre-med/pre-health oriented undergraduate population. The discussion-based format of the course was designed to generate dialog between lifelong learners and undergraduate students.

The course was formatted specifically to encourage discussion among individuals with different scientific backgrounds. Because many lifelong learners had little previous exposure to science, the course included coverage of extensive biography and social issues as well as background on the scientific and neurological principles described in the writing of Oliver Sacks. This text provided a unique combination of biography and neuroscience and was written so that it could be understood by those with a limited scientific background while providing a platform for the learning of fundamental principles of neuroscience and neurological disease. In addition, students were assigned small discussion groups that rotated each class and were designed to mix undergraduate and lifelong learning students. All class-wide discussions were first discussed in small group settings, with the aim of encouraging interactions among students.

Each class began with a discussion of a reading from Oliver Sacks’ autobiography ‘On the Move: A life’ [10]. This reading provided context for understanding Dr. Sacks’ unique approach to patients and medicine. It also provided a context to discuss social issues, including how one chooses a profession, friendships and family, changing views of sexuality, and how society handles sickness. These readings regularly generated lively discussion among undergraduate and Lifelong Learning students about historical differences (e.g., drug use in the 1960’s). There were also many discussions about the unique character and intellect of Dr. Sacks and how this contributed to his creative approaches to medicine.

Following the discussion of the autobiography, class meetings included discussion of multiple case studies from Oliver Sacks publications. The case studies were selected to cover a broad range of neurological issues ranging from development to neurodegeneration, providing a clinically relevant platform to introduce basic concepts in neurobiology and medicine [11–14]. The instructor provided a brief (∼10 minute) lecture-style introduction to each case study, focusing on the underlying biological concepts for those students in the class without a background in biology or neuroscience. The discussion of each topic was then facilitated by an undergraduate student. Round table discussion often focused on topics ranging from the mechanisms of the disorder under discussion to the physician-patient relationship.

## Method

### Participants

The Oliver Sacks course was made available to undergraduate students through standard mechanisms that included the posting of a brief description on the campus website and notification of advisors. To avoid selection bias among the undergraduate students, the inclusion of lifelong learners in the course was not in the course description. The course was classified as an upper division elective for Biology and Neuroscience Majors and was open to students in the College of Science and Wilkes Honors College of FAU. The course filled to capacity prior to the first day of classes. Lifelong learners were recruited through email and direct contact by Ms, Kristen Robbins, the coordinator of Jupiter’s Osher LLI. Six Lifelong Learners signed up for the course, with a commitment to regularly attend the class throughout the semester.

One lifelong learning student was unable to complete the class due to a health issue, and two undergraduate students joined the class at a later time and therefore are not included.

Seventeen students completed both the pre and post questionnaire. Participants were 5 lifelong learning students age 73 to 85 (*M*age = 77.60, *SD* = 5.08), six College of Science undergraduate students age 21 to 23 (*M*age = 22.00, *SD* = .89), and six Honors College undergraduate students age 20 to 22 (*M*age = 21.17, *SD* = .75). For the purposes of analysis, we have combined the undergraduate students. All questionnaires and procedures were approved by the Florida Atlantic University Institutional Review Board.

### Procedure

Both undergraduate and lifelong learning students completed a survey on the first day of class and another survey on the last day of class. The surveys were coded to protect the identities of the participants and included questions about participants’ expectations for the course, views on intergenerational learning, and educational background.

## Results

### Frequency of Intergenerational Interaction

The intergenerational class was designed to encourage interactions among younger undergraduate students and older lifelong learning students. Before taking the class, the undergraduate students reported varying levels of interaction with adults who were substantially older or younger than themselves. On the first day of class, students were asked to respond to the statement, “I regularly interact (on a weekly basis) with adults of a different generation.” on a scale of: 1 = strongly disagree, 2 = disagree, 3 = neutral, 4 = agree, and 5 = strongly agree. The majority of the students agreed (4 undergraduate and 2 lifelong learning) or strongly agreed (5 undergraduate and 1 lifelong learning) that they interacted on a weekly basis with adults of a different generation. One undergraduate selected neutral, and four students did not interact regularly with adults of a different age (2 undergraduate and 2 lifelong learning). Over the course of the term, the students all interacted with adults of different ages weekly in class.

### Value of Intergenerational Interaction

All the students valued the interactions they had with students of the other generation. At the end of the term, students strongly agreed (10 undergraduate and 4 lifelong learner) or agreed (2 undergraduate and 1 lifelong learner) with the statement, “I valued interactions with students from a different generation.”

Almost all of the students were enthusiastic about being a part of an intergenerational class even though the undergraduate students did not know the nature of the course before they enrolled. On the first day of the course, 82% of students (10 undergraduate and 4 lifelong learning) strongly agreed that “individuals of varying ages bring experiences and knowledge to the classroom that benefit my own learning,” and 12% (1 undergraduate and 1 lifelong learning) agreed with this statement. Only one undergraduate student strongly disagreed. On the final day of class, all of the students agreed (N = 5) or strongly agreed (N = 12) that having people of different ages enhanced their own learning. Thus, all of the students thought they benefited from the intergenerational classroom experience. At the end of the course, almost all of the students strongly agreed (11 undergraduate and 2 lifelong learning) or agreed (3 lifelong learning) that they were “likely to take a future class in this format.” Only one undergraduate student reported being neutral and no students disagreed.

### Comfort with Class Discussion

We hypothesized that the intergenerational format of the class would enhance students’ comfort with participating in class discussions. Students began the class with varying levels of comfort with class discussions. Among the undergraduate students, 6 of the students strongly agreed that they were comfortable participating, 2 agreed, 1 was neutral and 2 disagreed. Among the lifelong learning adults, 3 strongly agreed and 2 agreed. By the end of the course, all of the students were at least neutral, 71% strongly agreed (8 undergraduates and 4 lifelong learners) and 12% agreed (3 undergraduates and 1 lifelong learner) that they were comfortable with class discussions. One undergraduate student was neutral. Although the sample size is small, it is encouraging to see that by the end of the course none of the students felt uncomfortable participating in class discussions.

At the beginning of the term, we asked students if they preferred lecture-based or discussion-based courses and why. Ten undergraduate students reported preferring discussion classes and gave explanations like, “they allow you to benefit from hearing the varied opinions and knowledge from those with a different background” and “rather than memorizing concepts and studying with the sole purpose of taking an exam, a discussion-based course allows me to actually contribute to the development of the course.” Four students preferred lecture-based courses, giving the following reasons, “anxiety,” “quite shy,” “I am not yet comfortable with speaking my ideas. I often become insecure with my thoughts and opinions,” and “faster learning, broader subject matter.” Three students had no preference.

On the first day of class, we asked students what their expectations for the intergenerational course were and did they have concerns with the intergenerational format. All of the lifelong learning students had positive expectations saying, “interesting to observe different age-related points of view,” “I want young students to be interested in neuroscience and also learn that older people are not mindless,” “learning new ideas,” “new ideas,” and “learning about ideas, values, and intellect of another generation.” Two of the students were concerned, one with “filling three hours with relevant discussion” and the other with asking “people to speak louder.”

All undergraduate students had positive expectations for the course and made statements like, “I expect to learn a lot of different views,” “I expect to be able to get better at discussing opinions with people of all ages,” “I expect to gain a lot of wisdom and knowledge,” and “I expect to learn more and broaden my view on issues discussed in this course. I hope to learn something new from every classmate.” Comments also included “I expect to hear many different insights and opinions, a mixed group is great because as we grow old our opinions and perceptions greatly change,” “I expect to participate in engaging conversations that will help enable me to think more critically,” “I expect a lot of diverse perspectives filled with new knowledge and wisdom,” and “it should be fascinating to have mixed aged students in a discussion based course. The various life experiences would allow for extremely diverse viewpoints.” One student expressed concern about their grade.

At the end of the term, we asked students about their experience, and all of the participants reported having a positive experience. The LLI students made comments like, “I was awed by the knowledge of the students, and yet they welcomed us ‘oldsters’ and seemed anxious to hear our experiences and points of view. It was a very rewarding experience, and one I have shared with other lifelong learners who are now interested in participating in other courses of this type if they have the opportunity,” and “It has been an honor to know these young students and increased my appreciation for FAU. “The undergraduate students made comments like, “each person has experienced something different so by taking this class you allowed others to give input and their opinion. Not everyone is the same in what they believe so I found this class super valuable,” and “I learned so much from others in this class, the way they think is so different from how I think. I love their critical analysis of each chapter.” The honors students made comments like, “I had no idea what to expect, however, I enjoyed it a lot. Being able to hear opinions and knowledge from different age groups was really interesting,” and “it was great listening to insights from all other students each with different past experiences. The students from LLI enhances the discussion exponentially. I really enjoyed the class.”

## Discussion

In many intergenerational education programs involving older adults, the older adults serve as mentors and/or tutors to children and/or younger adults[15,16]. This is based on the premise that senior adults have experienced a career, while undergraduate students are preparing for the workforce. Older adults, however, can also learn from interactions with younger adults, and younger adults may learn more from older adults when those younger adults have more to contribute to the relationship and thus are more equal partners in [17]. Our findings support the premise that adults in each group have different strengths, knowledge, and life experiences that will be mutually beneficial. The mutual enthusiasm for the intergenerational classroom suggests both lifelong learners and undergraduates are engage one another as peers under these conditions.

Lifelong learning programs provide an alternative to degree-oriented programs and represent a primary form of educational access for older adults and retirees. Participation in these programs has profound effects on participants and local communities. For example, participation in informal learning has been shown to promote psychological and physical health [18,19]. Informal learning has potential to bring older adults together and provide a forum for social interactions and networking. Finally, lifelong learning has been shown to promote community and political engagement, extending the benefits of individual lifelong learning to the larger society. While these programs are widely viewed as successful, the curriculum typically lacks extensive science programming, perhaps because it is viewed as less accessible than arts, literature, and social science. While most lifelong learners had limited background in life science, the discussion-based format of the course allowed them to be active participants. Given that a limited number of upper-division elective courses is currently a constraint in accommodating the growth of pre-med/pre-health majors across the country, the generation of similar discussion-based classes ay meet the needs of lifelong learners and undergraduates.

Lifelong learning programs are increasing in prevalence but often lack interactive formats for enhancing science literacy in seniors [20]. While these programs have been remarkably successful, STEM education is underrepresented in educational programing. For example, our analysis at Florida Atlantic University, the nation’s largest lifelong learning program, fewer than 5% of classes were STEM related in 2016. This likely reflects limited ability to recruit STEM educators or lack of effective methodology for conveying principles in STEM to older individuals. Because science changes rapidly, it is important to provide STEM learning opportunities to lifelong learners with extensive, as well as limited, background in STEM.. The paucity of STEM access for lifelong learners highlights the need for novel mechanisms for inclusion of STEM-based classes in lifelong learning programs and more generally for improved access to STEM education for adult learners.

A limitation of most lifelong learning programs is that the members tend to be relatively homogeneous in age and ethnicity, preventing diverse interactions that are widely viewed as beneficial to both the individual and society. While the benefits of these programs have been well studied [21,22], lifelong learning programs typically generate interactions within a given peer-group. While FAU’s Osher Lifelong Learning Institute does not collect demographic data, the community is predominantly white with an estimated average age of over 70 (Robbins & Watlington, unpublished observations). By contrast, 89% of FAU undergraduate students are under the age of 35 and 37% are between 18 and 22. FAU has been nationally recognized as an ‘Opportunity College and University’ by the Carnegie Classification for its strong commitment to student success, particularly in expanding access for Pell Grant recipients and students from historically underrepresented backgrounds, and supporting their positive outcomes after graduation. In addition to enhancing STEM education, intergenerational classrooms may have additional positive benefits. We have known for many years that social support helps older and younger adults deal with difficult events in their lives e.g[23] and that older adults tend to have smaller social networks than younger adults [24]. The intergenerational interactions encouraged by the proposed STEM educational opportunities likely increase the social interactions of lifelong learners and undergraduate students. The undergraduate students may also benefit from interactions with the lifelong learners because older adults have been found to choose better strategies for success in dealing with interpersonal problems [25]. Furthermore, the classroom experiences led to intergenerational interactions that extended beyond the classroom, including a number of retirees working in research laboratories. Therefore, our results suggest an intergenerational classroom provides opportunities for adults of different ages to work together to enhance STEM learning and gain a greater understanding of, and learn from, people who are older or younger than themselves.

Undergraduate institutions can provide the infrastructure needed for teaching older adults, but this has been underutilized, despite the close proximity of undergraduate institutions and lifelong learning societies across the country. This infrastructure includes STEM faculty with diverse expertise in research and classroom settings who can offer a broad range of STEM courses, including interactive special courses, and lab experiences for students with limited science background. While most Lifelong Learning Programs are directly affiliated with colleges and universities and/or are located on university/college campuses [26–28] the academic programming for lifelong learners is usually separate. Providing lifelong learners access to undergraduate STEM courses or generating curriculum that encourage lifelong learners would greatly expand STEM access to this portion of the population and would generate mutually beneficial interactions between lifelong learners and undergraduate students. Integrating older adults into the undergraduate curriculum would generate opportunities for science literacy and promote intergenerational interactions.

